# Single-Cell RNAseq analysis of infiltrating neoplastic cells at the migrating front of human glioblastoma

**DOI:** 10.1101/165811

**Authors:** Spyros Darmanis, Steven A. Sloan, Derek Croote, Marco Mignardi, Sophia Chernikova, Peyman Samghababi, Ye Zhang, Norma Neff, Mark Kowarsky, Christine Caneda, Gordon Li, Steven Chang, Ian David Connolly, Yingmei Li, Ben Barres, Melanie Hayden Gephart, Stephen R. Quake

## Abstract

Glioblastoma is the most common primary brain cancer in adults and is notoriously difficult to treat due to its diffuse nature. We performed single-cell RNAseq on 3589 cells in a cohort of four patients. We obtained cells from the tumor core as well as surrounding peripheral tissue. Our analysis revealed cellular variation in the tumor’s genome and transcriptome, We were able to identify infiltrating neoplastic cells in regions peripheral to the core lesions. Despite the existence of significant heterogeneity among neoplastic cells, we found that infiltrating GBM cells share a consistent gene signature between patients, suggesting a common mechanism of infiltration. Additionally, in investigating the immunological response to the tumors, we found transcriptionally distinct myeloid cell populations residing in the tumor core and the surrounding peritumoral space. Our data provide a detailed dissection of GBM cell types, revealing an abundance of novel information about tumor formation and migration.

## Introduction

Glioblastoma (GBM) is the most common malignant primary brain cancer in adults (Bush et al., 2016). GBMs are incurable tumors; despite aggressive treatment including surgical resection, chemotherapy and radiotherapy, the median overall survival remains only 12-18 months (Wen and Kesari, 2008). Unlike brain metastases, for which local control rates following surgery and radiation can reach 80%, GBMs are diffusely infiltrating (Claes et al., 2007) and invariably recur, even in distant regions of the brain. The diffuse nature of GBMs renders local therapies ineffective as migrating cells outside of the tumor core are generally unaffected by local treatments and are responsible for the universal recurrence of GBMs in patients.

The development of novel treatment strategies is predicated upon a better understanding of the molecular features of these tumors, with a particular focus on the ability to capture and identify the infiltrating cells responsible for recurrence. Although bulk tumor sequencing approaches have been useful in generating classification schemas of GBM subtypes (Cancer Genome Atlas Research Network, 2008; Verhaak et al., 2010), they provide limited insight to the true heterogeneity of GBM tumors. Inter-patient variation and molecular diversity of neoplastic cells within individual GBMs has been previously described (Patel et al., 2014), but studies thus far have been limited in scope to the molecular complexity of cells from the tumor core; existing studies at single-cell resolution have been unable to address the nature of infiltrating GBM cells or the unexplored variety of other neuronal, glial, immune, and vascular cell types that reside within and around GBMs. The interplay between each of these cell types within the tumor microenvironment likely contributes significantly to tumor progression and resistance to therapy.

To capture and characterize infiltrating tumor cells, and to define the cellular diversity within both the tumor core and surrounding brain, we performed high depth single-cell RNAseq on a cohort of four primary GBM patients (IDH1 negative, Grade IV GBMs confirmed via pathological examination). From each patient, we collected samples from two separate locations: the first residing within the tumor core and the second from peritumoral brain (**Fig. 1A** and **Fig. S15**). Additionally, from each location we collected both unpurified cell populations, as well as populations enriched for each of the major CNS cell types (neurons, astrocytes, myeloid cells, endothelia) that are often overwhelmed in number by the abundance of tumor cells. This strategy allowed us to capture tumor cells that had migrated away from the primary tumor lesion into the peritumoral tissue, and to specifically compare transcriptome-level effects of the tumor microenvironment on each of the various brain and immune cell types.

**Figure 1.**
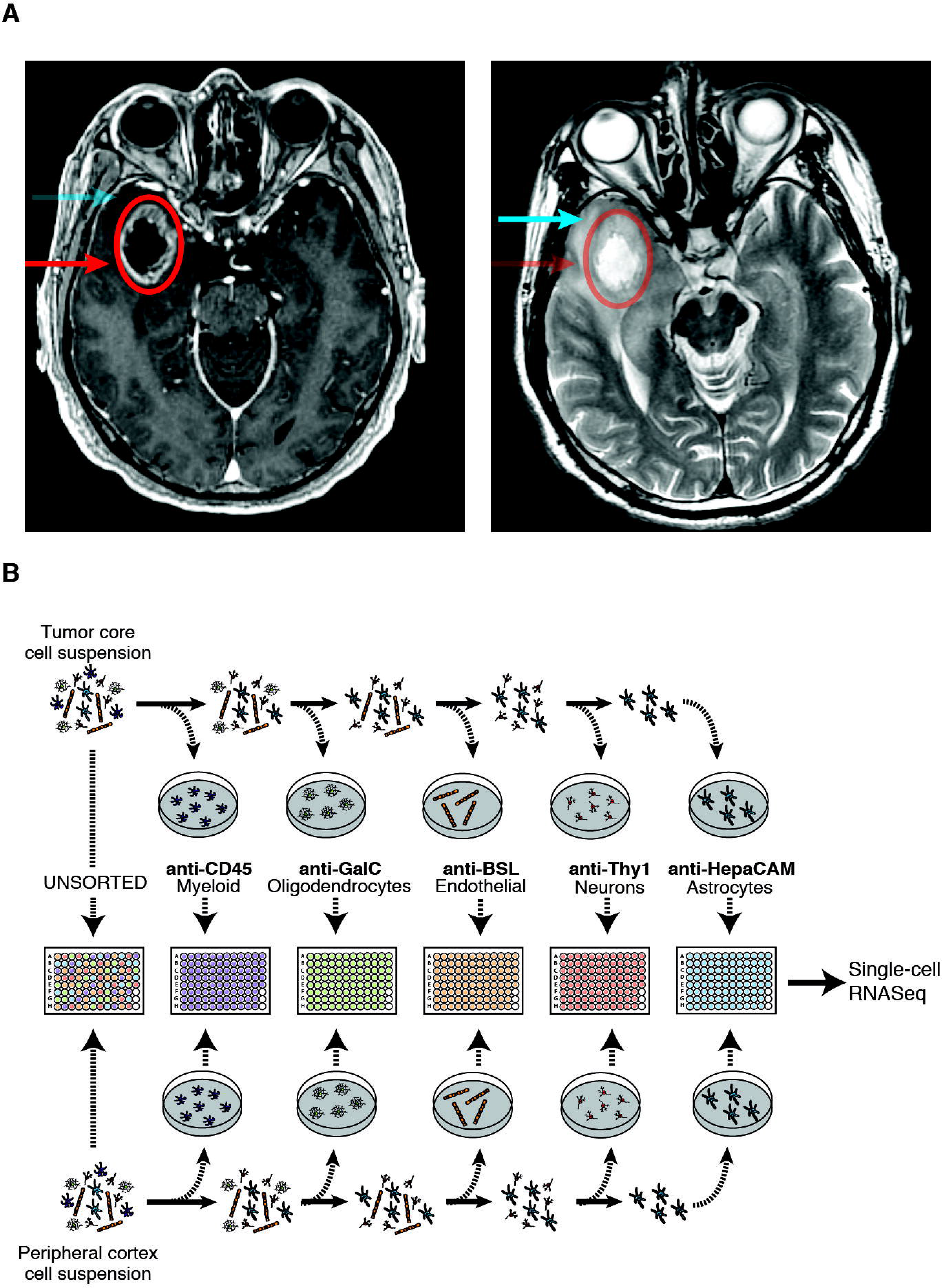
Experimental layout. (A) Axial T1 with contrast (left side) and T2 (right side) MRI brain in a patient with a right temporal GBM. The tumor core was defined as contrast enhancing (red circle, arrow), and the peritumor brain was non-contrast enhancing, yet T2 hyperintense (blue arrow). (B) Overview of the experimental procedure.

In total, we sequenced 3589 cells from both the tumor core and the peritumoral brain, comprising neoplastic cells and representatives from each of the major CNS cell types (vascular, immune, neuronal, glial). Together, our data provide a large-scale dissection of GBM cell types and their respective gene expression profiles, revealing an abundance of information about tumor formation and effects of the interaction between tumor cells and the immune system. Specifically, we managed for the first time in primary tumors to capture and characterize infiltrating neoplastic cells along the migrating front of the GBM. Additionally, we investigated the heterogeneity of GBM tumors between and within patients, and characterized the effect of the tumor environment on populations of non-neoplastic cells with particular emphasis on immune cell populations.

## Results and Discussion

### Initial clustering and identification of cell types

At the onset of our efforts to sort single cells from the tumor core, we discovered that the vast majority of the cells we captured from dissociated tumor belonged to the neoplastic population, with little contribution from other glial, neuronal, vascular, or immune subtypes. Thus, to increase the relative percentage of non-neoplastic cells in our analysis, we adapted well-validated protocols for immunopanning human tissue with cell-type specific markers (Zhang et al., 2016) with the ultimate goal of encompassing the entirety of the tumor and peritumor cellular landscape that is often blurred in bulk sequencing studies or insufficiently sampled in prior single cell studies. After dissociation and immunopanning, individual cells were sorted into 96-well plates; non-immunopanned cells were also sorted to ensure that no major subpopulation of cells was excluded. The details of this process is described in detail in the Materials and Methods and graphically summarized in Fig. 1B.

To visualize the transcriptomic landscape across all sequenced single cells, we used dimensional reduction to generate a two dimensional map of all 3589 single cells that passed QC (Fig. S1 and Table 1), performing an analysis similar to Darmanis et al (Darmanis et al., 2015). Briefly, we selected genes with the highest over-dispersion (n=500) and used them to construct a cell-to-cell dissimilarity matrix. We then performed t-Distributed Stochastic Neighbor Embedding (tSNE) on the resulting distance matrix to create a two-dimensional map of all cells. Finally, we used k-means clustering on the 2D tSNE map, resulting in the identification of 12 distinct cell types within separate clusters (Fig. 2A).

**Figure 2.**
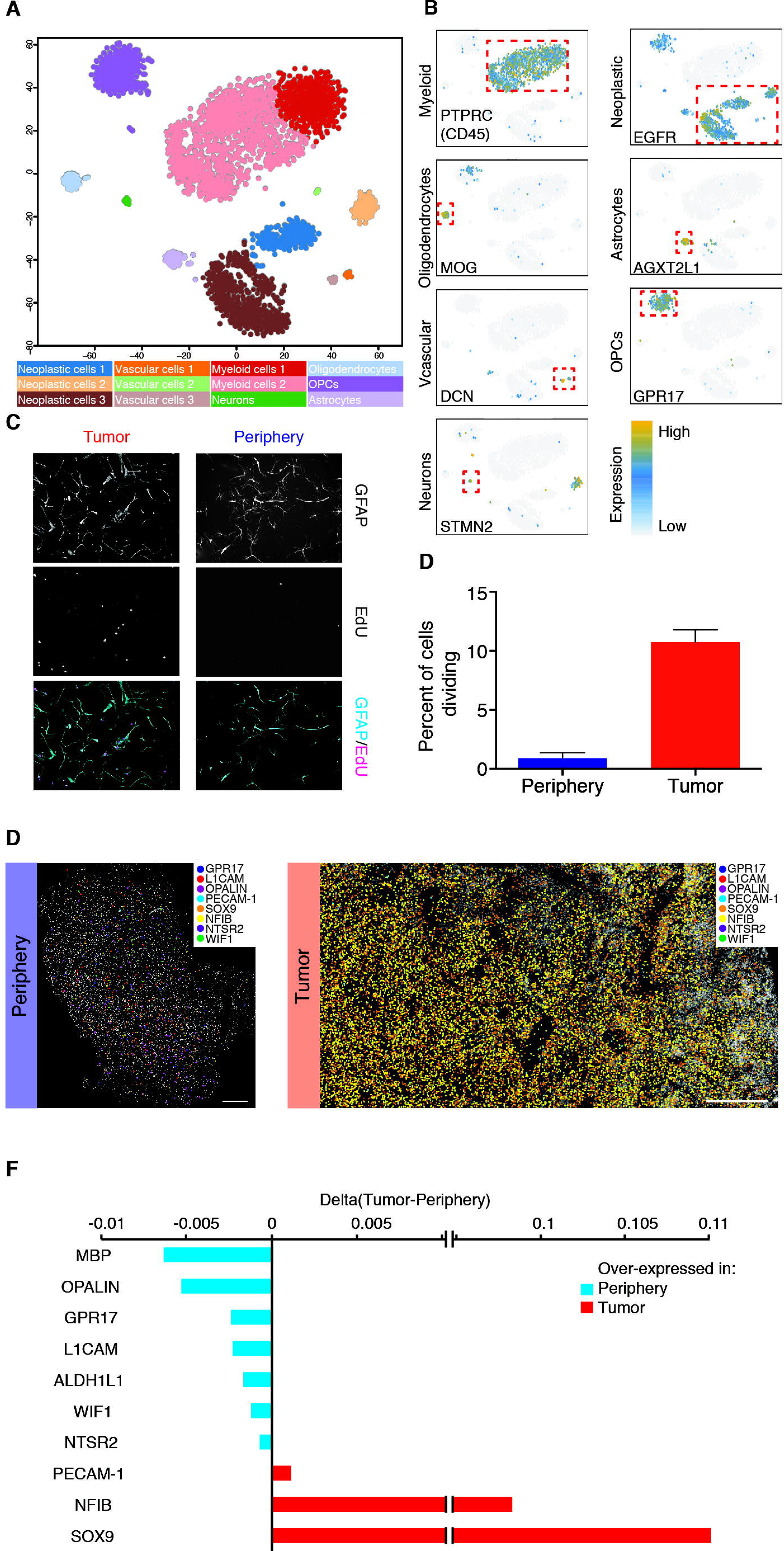
General characteristics of neoplastic cells. (A) 2D-tSNE representation of all single cells included in the study (n=3589). Cell clusters are differentially colored and identified as distinct cell classes. (B) Expression of characteristic cell type-specific genes overlaid on the 2D-tSNE space. (C) GFAP and EdU staining of HEPACAM selected cells. (D) Quantification of EdU positive cells as a percentage of total DAPI nuclei from the tumor core and the surrounding peritumor brain in culture for 7 days. (E) *in situ* RNA staining of neoplastic and non-neoplastic specific genes in tumor tissue (right) and peritumoral brain (left). (F) Quantification of *in situ* RNA signals shown in Figure 2E. Difference between tumor and peritumoral brain is shown.

**Table 1.** Dataset summary. Number of cells per sample, number of cells per anatomical location and general sequencing statistics summarized for all sequenced cells that passed or failed QC. Only cells that passed QC were further analyzed.

|  | Pass QC | Fail QC |
| --- | --- | --- |
| Cells | 3589 (89%) | 456 (11%) |
| Sample |  |  |
| BT_S1 | 489 | 11 |
| BT_S2 | 1169 | 138 |
| BT_S4 | 1542 | 267 |
| BT_S6 | 389 | 40 |
| Location |  |  |
| Tumor | 2343 | 212 |
| Periphery | 1246 | 227 |
| Sequencing statistics |  |  |
| Reads (median) | 1706000 | 1019000 |
| Uniquely mapped reads % (median) | 75.81 | 81.93 |
| Multimapped reads % (median) | 2.35 | 0.29 |
| Genes detected (median) | 2129 | 33 |

We inferred the cellular identities of the resulting clusters from the tSNE analysis by identifying significantly overexpressed genes in each cluster. Initially, we used a Wilcoxon rank sum test to compare each gene’s expression in one cluster to its expression in all other cells, and we then cross-referenced these cluster-enriched genes (Table S1) with known cell-type specific CNS databases (Zhang et al., 2016; 2014). We then validated the identity of the clusters by combining our dataset with published single cell RNAseq data from healthy human brain samples to ensure consistent cell type classifications of each cluster (Darmanis et al., 2015). The cell type identity of each cell cluster along with the number of cells originating from each patient and anatomical location are shown in Table S2. Genes whose expression is specific to major cell types of the brain are shown in Fig. 2B. Using this method, we were able to classify all clusters into one of the major CNS neuronal, glial, or vascular subtypes found within the healthy human brain, with the exception of three clusters (1, 4 and 11). After further investigation, we preliminarily defined the cells within these three clusters as ‘neoplastic’ based on the following observations (as well as CNV analyses performed below). Foremost, ~94% of the cells (1029/1091) in neoplastic clusters originated from the tumor core. Additionally, these cells significantly over-expressed EGFR (Fig. 2B), a gene upregulated in 30-50% of all GBMs (Libermann et al., 1985a; 1985b; 1984), as well as SOX9, a transcription factor with an established oncogenic role in gliomas (Wang et al., 2012a). Interestingly, the combined expression of just these two genes, EGFR and SOX9, was capable of demarcating neoplastic cells with high sensitivity and specificity (Fig. S2).

To further increase our confidence in the identity of the inferred cell types (both typical and neoplastic), we performed a direct comparison with single cell and bulk RNAseq data from (Darmanis et al., 2015) (Healthy Brain) and (Patel et al., 2014) (Bulk GBM). The Healthy Brain dataset contains single cell RNAseq data from 332 cells originating from healthy adult human cortex and the Bulk GBM dataset contains RNAseq data from bulk sequencing of five primary GBMs. The resulting tSNE map of all cells and bulk samples can be seen in **Fig. S3**. Using this extended dataset we made two important observations; Bulk GBM samples cluster directly in the midst of our neoplastic cell clusters, while single cells from the Healthy Brain dataset, originating from tumor free brains, cluster together with non-neoplastic cells in each of their respective assigned cell types (**Table S2**). These observations provided further validation of our classification schema for both the identity of non-neoplastic cell types as well as the identification of GBM-specific neoplastic populations.

**Figure 3.**
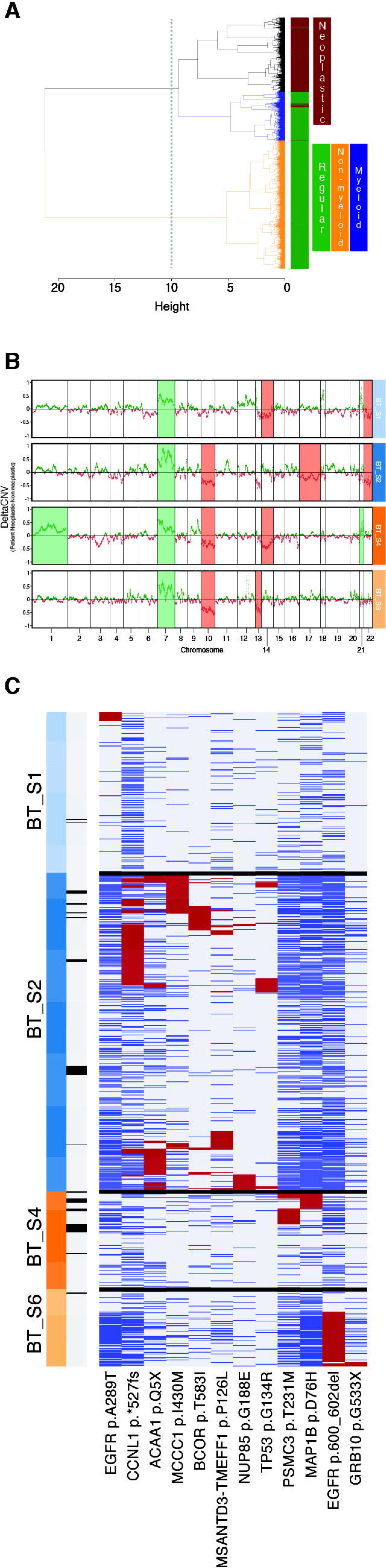
(A) Hierarchical clustering of all cells on the basis of their RNA-seq derived CNV profile. Dendrogram branches are colored to denote different clusters of cells. Color bar demarcates neoplastic (brown) vs non-neoplastic (green) cells. (B) Delta_CNV_ profils of each patient’s neoplastic cells and non-neoplastic non-myeloid cells. Specific chromosomes that were found to be over- or under-represented in each of the patients are highlighted. (C) Exonic non-synonymous variants occurring in greater than 5% of any patient’s neoplastic cells. Cells (columns) may either contain a variant (red), not contain the wildtype (blue), or display insufficient read coverage at that position to make a determination (gray). Cells are labeled by patient of origin (top color bar) and infiltrating status (bottom color bar; dark is infiltrating)

Since the majority of cells that were classified in the neoplastic clusters originated from the tumor core and were selected by HepaCAM immunopanning (an astrocyte-lineage marker), we maintained some of these cells in culture to compare morphological and proliferative features to HepaCAM selected cells from the surrounding tissue. Compared with peripheral astrocytes, HepaCAM-selected cells from the tumor core exhibited a distinct morphology (Fig.2C) and were highly proliferative, as determined by EdU incorporation and expression of *MKI67* (Fig.2D), further supporting the notion that they are neoplastic in origin.

Although the tumor core and surround samples were separated by MRI-guided surgical excision, we also looked for a molecular signature to help confirm the identify of the tumor core cells from the surrounding cells. We hypothesized that expression of hypoxic genes would correlate with the relatively oxygen-poor tumor core where cells are forced to adapt to the low-oxygen environment. Thus, we calculated a ‘hypoxic-score’ for each cell based on the combined expression of classic hypoxic genes *PGK1*, *CA9, VEGFA, SPP1*, and *HIF1A* (**Fig. S4**). As expected, we observed a significant increase in the expression of these hypoxic genes within the tumor core as compared to the surrounding regions.

**Figure 4.**
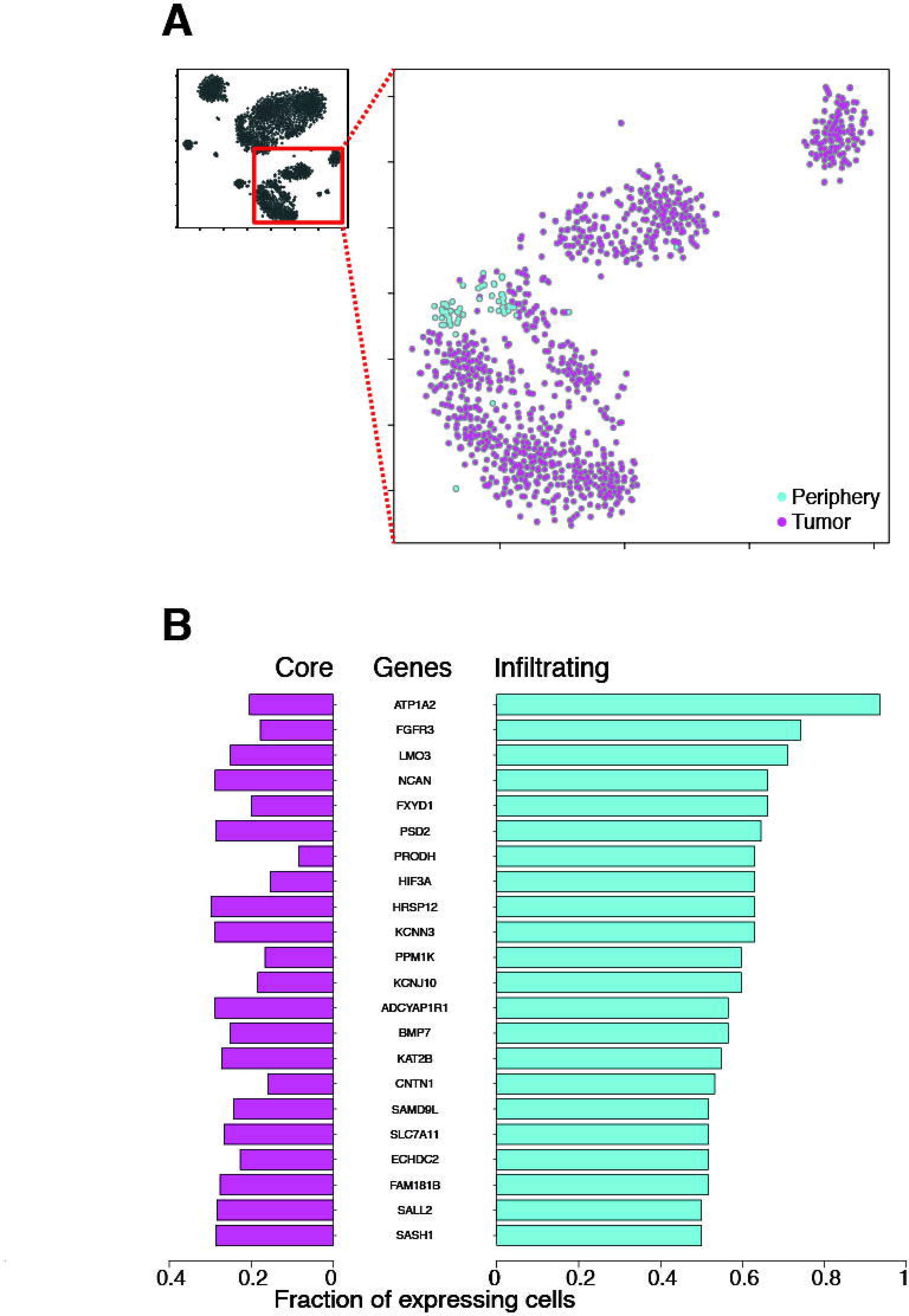
Analysis of infiltrating tumor cells. (A) 2D-tSNE representation of all neoplastic cells colored by location (tumor vs periphery). (B) Differentially expressed genes between neoplastic cells originating from the tumor (Core) or the periphery (Infiltrating). The fraction of tumor core and infiltrating cells expressing any given gene is plotted.

It should be noted that the tumors themselves are heterogeneous and contain both neoplastic cells and non-neoplastic cells. Out of 2343 tumor core-originating cells, only 1029 were members of the aforementioned neoplastic-cell clusters (~44%). The vast majority of the remaining cells (n=1182, ~50%) belonged to immune cell clusters, with the residual cells assigned to the OPC cluster (n=50, 2.13%), one of the endothelial cell clusters (n=47, 2%), the oligodendrocyte cluster (n=34, ~1.5%), or the neuronal cluster (n=1, ~0.05%). Interestingly, the only cell population without a cell originating from the tumor tissue sample is the mature astrocyte cluster.

### Neoplastic cell characteristics

The neoplastic cells share common characteristics that distinguish them from other cells of the brain. Differential expression analysis (DESeq2) between neoplastic and non-neoplastic cells revealed genes enriched within all neoplastic cells (**Fig. S5A**, also see Supplementary information). Unsurprisingly, *EGFR* showed the highest enrichment of any gene. In addition, among the highest enriched genes we found *CHL1*, a member of the L1-family of neural cell adhesion molecules that is involved in migration and positioning of neurons in the developing neocortex. Furthermore, we found upregulation of the transcription factors *SOX2* and *SOX9*, which are also involved in brain development and lineage specification. Notably, *SOX2* has been reported as a marker of glioma stem cells, along with genes *POU3F2*, *OLIG2* and *SALL2* (Suvà et al., 2014), while *SOX9* has been shown to correlate with poor clinical outcome (Wang et al., 2012b). *NFIB* was another neoplastic-specific transcription factor that has been implicated in brain development (Campbell et al., 2008; Steele-Perkins et al., 2005) with an important role in the induction of quiescence in neural stem cells (Martynoga et al., 2013) and promotion of metastasis (Denny et al., 2016). We verified the upregulation of both *NFIB* and *SOX9* using multiplexed *in situ* RNA staining, with padlock probes and rolling circle amplification (Ke et al., 2013), on sections of tumor core as well as peripheral tissue (**Fig. 2D and Fig. S6**). We also looked for gene expression that was specifically limited to non-neoplastic cell populations and verified their expression patterns via *in situ* RNA staining (**Fig. 2D and Fig. S6**). Unsurprisingly, many of these markers were genes known to define mature, differentiated CNS cell types: MBP and OPALIN (oligodendrocytes), GPR17 (OPCs), L1CAM (Neurons), as well as ALDH1L1, WIF1 and NTSR2 (Astrocytes).

**Figure 5.**
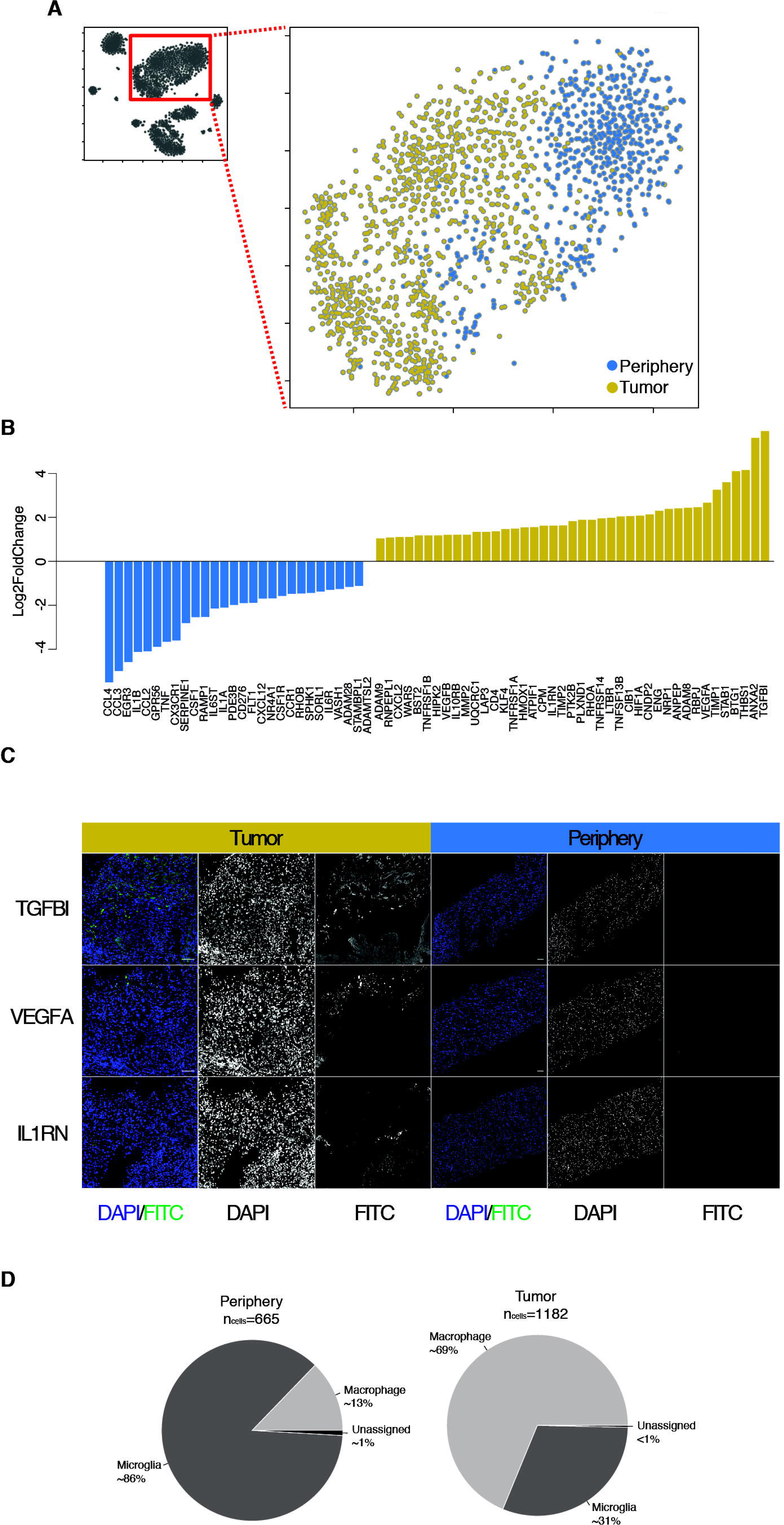
Analysis of immune cells. (A) 2D-tSNE representation of all immune cells colored by location (tumor vs periphery). (B) Barplots of Log2 fold changes between immune cells from the tumor and periphery for a curated list of genes involved in ECM remodeling, angiogenesis and immune regulation. (C) IFC stanining of TGFBI, VEGFA and IL1RN in tumor (left) and peripheral tissue (right). (D) Percentage of immune cells from the tumor or periphery classified as macrophages or microglia

In light of recent advances in our understanding of how tumors affect immune cells within their microenvironment, as well as a number of clinical studies pursuing the use of immune checkpoint inhibitors for the treatment of GBM, we next looked at the expression of MHC I genes **(Fig. S7A)** and genes coding for ligands of PD1 (*CD274* and *PDCD1LG2*) and CTLA4 (*CD80* and *CD86*) on neoplastic cells **(Fig. S7B)**. We found that most neoplastic cells do not express the transcripts for ligands of *PD1* and *CTLA4*, whereas the opposite is true of genes of the MHC I class. Nonetheless, we did notice a high degree of heterogeneity both within and between patients. Specifically, fewer than 25% of neoplastic cells of BT_S1 express *HLA-A, HLA-B*, and *HLA-C*, while more than 75% of neoplastic cells from patients BT_S2, BT_S4 and BT_S6 express these genes **(Table S4)**. We observed the same heterogeneity in the expression of ligands of PD1 and CTLA4 as well. Interestingly, most of the neoplastic cells do not express CD274, PDCD1LG2, CD80 or CD86 with the exception of a small subset cells from BT_S2 expressing both ligands of PD1 **(Fig. S7C)**, which suggests that therapeutics directed against these targets (so-called checkpoint inhibitors) may have limited efficacy in this tumor type. We also compared the expression of immune checkpoint receptor ligands in myeloid cells from the tumor core and the peritumoral tissue, as discussed below.

Within the greater neoplastic cell cluster we found a high degree of both inter- and intra-tumor variation. From an inter-tumor perspective, we observed that neoplastic cells largely separated based on patient of origin with each patient’s neoplastic cells clustering preferentially with other neoplastic cells from the same patient (**Fig. S8A**); this enabled identification of genes that differentiated between each patient’s tumor (**Fig. S8B**). Inter-patient heterogeneity was even more apparent at the level of unique CNV abnormalities between patients, as discussed below. From an intra-tumor perspective, we wondered whether heterogeneity within individual patient tumors may reflect differences in the Verhaak classification (Verhaak et al., 2010) of these tumors. However, like previous findings (Patel et al., 2014), we found that each patient’s tumor represents an ensemble of cells belonging to each of the different Verhaak molecular subtypes (see Supplementary Information). We therefore quantified the degree of intra-tumor heterogeneity, and used the endogenous heterogeneity as other CNS cell types as a benchmark for comparison (i.e. oligodendrocytes, astrocytes, OPCs). In **Supplementary figure 6C**, we plotted histograms of the distributions of all pairwise distances (self-to-self distances were removed) between neoplastic cells of each patient and cells belonging to each the non-neoplastic clusters. We observed a clear and statistically significant (p < 10^-16^) difference between the distribution of distances of neoplastic cells originating from each of the patients and those found within the healthy cell clusters, suggesting that the neoplastic cell clusters had a higher level of internal transcriptomic heterogeneity compared to non-neoplastic CNS cell types.

### Neoplastic cell CNV analysis using RNAseq

Genomic Copy Number Variants (CNVs) are known to be among the triggers of tumor formation (Shlien and Malkin, 2009), and tumor progression is commonly associated with further variations in copy number. Using the RNAseq data, we calculated CNV vectors for each individual cell and then clustered cells on the basis of their respective profile CNV vector (as opposed to the typical gene expression profile used for tSNE analysis). The resulting dendrogram was comprised of three primary branches (**Fig. 3A**): one (CNV 1) consisted exclusively of neoplastic cells, while the remaining two contain the majority of non-neoplastic cells. More specifically, one of the two non-neoplastic clusters contained all CD45^positive^ antigen-presenting (AP) cells (CNV 3), and the remaining branch consisted of all other non-neoplastic cells (CNV 2). The most likely reason for the separate clustering of normal and AP cells is the overexpression of the MHC class II genes, which cluster together on chromosome 6 (data not shown). To evaluate the degree of correlation between the CNV analysis and the initial tSNE clustering, we calculated the agreement between classifications in the two platforms. Of the cells identified as neoplastic in the initial tSNE plot, 1047/1091 were similarly identified as neoplastic in CNV1 (4% misclassification). Additionally, 2483/2498 cells that were initially identified as non-neoplastic belonged to either of the non-neoplastic clusters (CNV2 or CNV3) (0.6% misclassification).

To determine which chromosomal regions were most affected in neoplastic cells, we subtracted the median CNV profile of the non-neoplastic, non-myeloid populations (CNV 2) from that of the neoplastic cells (CNV 1). The individual median CNV profiles along with their differences (Delta_CNV_) can be seen in **Fig. S9A**. If the differences between RNAseq-derived CNV profiles across different cell populations were merely a result of gene-expression fluctuations with no underlying structural chromosomal variation, we would expect the Delta_CNV_ in Fig. S9A to randomly fluctuate around zero. Instead, we specifically looked for the presence of entire chromosomes that were over or under-represented in the neoplastic cells by counting the number of positive (higher in neoplastic cells) and negative (higher in non-neoplastic cells) occurrences (**Fig. S9B**) for each chromosome. The RNA-derived CNV profiles of neoplastic cells revealed two well-described chromosomal alterations in gliomas (Reifenberger and Collins, 2004): the amplification of chromosome 7 (including *EGFR*) as well as a putative deletion of chromosome 10 (including *PTEN*, *MGMT*). In addition, we noticed a similar pattern for chromosome 22 as we did with chromosome 10, suggesting another GBM specific putative deletion. While the amplification of chr7 is observed in all four patients, inter-patient heterogeneity is largely reflected in the neoplastic CNV profiles for each of the patients’ tumors, as can be seen in **Fig. 3B**.

To validate the utility of the RNA-derived CNV profiles, we performed DNAseq on DNA isolated from the tumor and the peritumoral tissue of one of the patients (BT_S4). We found that RNA-derived CNV profiles closely represented those derived from genomic data. As predicted, bulk DNAseq revealed a richer landscape of chromosomal aberrations in the tumor, not evident in single-cell-RNAseq-derived CNV profiles. Nonetheless, all identified chromosomal aberrations (over-representation of chromosomes 1, 7 and 21 and under-representation of chromosomes 10 and 14) in that patient’s tumor found by RNAseq were similarly classified by DNAseq (**Fig. S10**).

### Single-cell genomic variant analysis using RNAseq data

Single-cell RNAseq data also have the capacity to reveal clinically relevant tumor variants, such as SNVs as well small indels, in heterogeneous tumors. We called variants using the GATK pipeline (see Variant analysis section in Supplementary information) for single cell RNAseq data and subsequently limited the variant set to non-synonymous exonic variants occurring in at least 5% of a patient’s neoplastic cells. To exclude likely germline mutations we removed variants observed in the 1000 Genomes Project, those found in greater than 3 non-neoplastic cells, and those where greater than 2% of non-neoplastic cells with reads covering the variant position expressed the variant allele. Cross-referencing the remaining putative tumor-specific variants with previously identified somatic tumor variants using the Catalogue of Somatic Mutations in Cancer led to the identification of neoplastic and patient specific subpopulations harboring both novel and well-established mutations on a number of tumor-related genes, absent in non-neoplastic cells (**Fig. S11**). Specifically, we found two EGFR variants, a previously reported missense substitution (c.G865A:p.A289T, COSM21686) and a novel non-frameshift deletion of nucleotides 1799 to 1804 on exon 15. Interestingly, the first variant is found on a subset of BT_S1 neoplastic cells and the second in a subset of neoplastic cells of BT_S6 (**Fig. 3C**). Neoplastic cells of patient BT_S4 harbor a missense mutation (c.G226C:p.D76H) on MAP1B, which is involved in regulation of cytoskeletal changes occurring during neurogenesis. We were able to confirm this variant as tumor-specific using whole genome sequencing of BT_S4 tumor and peripheral bulk DNA. Of note, the same position of MAP1B has been previously found mutated in skin carcinomas but with a different substitution (c.G226A:p.D76N, COSM5908518). In a subpopulation of BT_S2 neoplastic cells we observed a potential cancer driver mutation in the form of a missense TP53 mutation (c.G226A:p.D76N, COSM5908518), previously found in multiple cancer types including gliomas as well as lung and breast carcinomas. In addition, a subset of cells from the same patient carry a CCNL1 frameshift mutation in exon 11 resulting in deletion of the stop codon as well as a missense mutation (c.C1748T:p.T583I) in BCOR, a gene which has previously been shown to be somatically mutated in glioblastoma (Frattini et al., 2013). Additional tumor-specific somatic tumor variants are illustrated in **Fig. 3C**, which further highlights intra-tumor heterogeneity and inter-patient heterogeneity.

### Glioblastoma infiltrating tumor cells

We observed that although the vast majority of cells (n = 1029) within the neoplastic clusters were collected from the tumor core, a small number of cells with a neoplastic signature originated from the peripheral tissue (n=62) (Fig. 4A). We hypothesized that these may represent neoplastic cells that migrated from the tumor core to the surrounding peritumoral space and hereafter refer to them as infiltrating cells. To further verify that these cells are neoplastic in origin, we found that 57 of 62 were found in cluster 1 (the neoplastic cluster) of the CNV analysis. Furthermore, the it is evident from the map of tumor-specific somatic variants in **Fig. 3C** that infiltrating cells are indeed neoplastic as they share a number of SNV variants with other core-derived neoplastic cells of each patient. A support vector machine (SVM) classification analysis using genomic variants, trained on non-infiltrating neoplastic and all non-neoplastic cells, also identified 57/62 cells as neoplastic in origin.

The infiltrating cells are unlikely to represent contamination from the tumor core since the peripheral tissue was the first sample to be removed during the surgery under MRI guidance. Furthermore, despite the inter-patient heterogeneity across neoplastic cells (each patient’s tumor cells form their own distinct clusters on the tSNE plot), the infiltrating cells from each patient clustered closely to each other irrespective of sample of origin. To better quantify the relationship between core and infiltrating cells, we used nearest-neighbor classification to predict the site of origin (tumor or peripheral) for each neoplastic cell based on the tissue of origin of three of its closest neighbors. Despite the fact that our dataset contains ~15 times more neoplastic cells from the tumor core compared to the periphery, we were able to correctly classify peripheral neoplastic cells for more than 85% of the cells. Unlike the cells residing in the tumor core, infiltrating cells seem to have down-regulated genes involved in adaptation to hypoxic environments as they are migrating through the (relative) oxygen-rich proximal brain tissue (p<10^-5^ lower hypoxic scores as determined by a Wilcoxon rank sum test) (**Fig. S4**).

To understand the molecular features that distinguish infiltrating cells, we performed differential expression analyses (DESEq2) and found ~1000 and ~250 genes that were down- or up-regulated, respectively, in infiltrating cells compared to tumor core cells. Further selection of differentially expressed genes expressed in greater than 50% of infiltrating cells and less than 30% of core neoplastic cells narrowed down the list of up-regulated genes to 22 (**Fig. 4B**). Among the top ten genes enriched in the infiltrating cell population we find genes with functions involving the invasion of the interstitial matrix. For example, size regulation may be achieved via the overexpression of Na+/K+-ATPases like *ATP1A2* (also further regulated by *FXYD1*) while the overexpression of enzymes like *PRODH* involved in proline catabolism may contribute to increased ATP energy demands of migrating cells. We also noticed increased infiltrating cell expression of cell survival signaling via the *FGFR3* receptor as well as *LMO3* via inhibition of *TP53* mediated apoptosis. FGF signaling-mediated survival has been previously linked to chemotherapy resistance. Interestingly, FGF signaling is important for cell migration during embryogenesis while re-activation of the pathway has been linked with cell migration and invasion in prostate tumors (Turner and Grose, 2010).

Gene Ontology analysis (**Fig. S12**) of genes upregulated in infiltrating cells revealed significant enrichment of GO categories (query space: Biological Processes) that are highly relevant to tumor cell migration (Demuth and Berens, 2004) such as cell-cell adhesion (*ECM2, ANGPT1, TSPAN7*), anion-transport (*TTYH1/2, AQP1*), nervous system development (*BCAN, HES6, GLI3*), as well as a number of metabolic processes. A similar analysis using PAGODA (Fan et al., 2016) gave consistent results (**Fig. S13**).

It has remained a longstanding question whether infiltrating GBM cells actively proliferate while disseminating throughout the brain. When we looked specifically at the proportion of proliferating cells within the tumor core or infiltrating neoplastic cells we found 7.7% (80/1029) of neoplastic tumor core cells were actively proliferating in comparison to only 1.6% (1/62) of infiltrating tumor cells, as determined by MKI67 expression. It should be also noted that within the cells included in this dataset, we noted the strong resemblance of neoplastic cells to OPCs (**Fig. S5B**) suggesting functional similarity or a possibly alternate cell-of-origin to astrocytes for GBM (Liu et al., 2011). Of note, we also observed a strong resemblance of neoplastic cells to fetal human astrocyte gene signatures from a separate dataset (Zhang et al., 2016).

### Analysis of immune cells

Two of the largest tSNE clusters that we observed were comprised of immune cells, defined by the specific expression of numerous myeloid-specific genes including *PTPRC* (CD45) as well as MHC class II genes (Fig. 2B). Further analysis, summarized in the Supplementary Information section, demonstrated that the vast majority of cells within these populations could be classified as either macrophages or microglia (>95%), with the remaining population comprised primarily of dendritic cells (~4.5%). Of the two primary myeloid clusters, we noticed that each contained cells almost exclusively from either the peritumoral space or tumor core, respectively (Table S2 and Fig. 5A), suggesting pronounced gene expression differences between the intra- and extra- tumor core myeloid subpopulations.

To better specify the identity of each myeloid cell within and surrounding the tumor core, we correlated gene expression of each cell with a panel of established macrophage and microglia specific genes (*TMEM119, P2RY12, GPR34, OLFML3, SLC2A5, SALL1*, *ADORA3* for microglia and *CRIP1, S100A8, S100A9, ANXA1*, *CD14* for macrophages) (Bennett et al., 2016) We then classified each cell as macrophage or microglia based on the combined expression of those genes, and found that the majority of cells within the tumor core tended to express genes characteristic of macrophages (n_macrophage_=813, n_microglia_=365) whereas cells from the surrounding space expressed genes characteristic of microglia (n_macrophage_=85, n_microglia_=574) (**Fig. 5D**). These data suggest that tumor-infiltrating macrophages and resident brain microglia preferentially occupy the tumor and peritumoral spaces, respectively.

We then examined the differential expression of selected gene sets within and surrounding the tumor core. These genes included cytokines, chemokines, chemokine receptors, matrix metalloproteinases (MMPs), and genes involved in angiogenesis. For a broader view of the role of tumor residing myeloid cells, we examined any gene of our curated gene list that was significantly (p_adj_<0.01, as determined by DESeq2) up or down-regulated in tumor or surrounding myeloid cells in at least one of the patients. We observed discrete gene expression differences between tumor and surrounding myeloid cells, with a predilection for pro-inflammatory markers expressed in the tumor periphery, and more anti-inflammatory / pro-angiogenic factors expressed in the tumor core (**Fig. 5B**). We subsequently validated the tumor core-specific presence of several biologically relevant examples of these molecules via *in situ* hybridization stainings. For example, while the inflammatory markers *IL1A/B* were up-regulated in the peritumor brain, *IL1RN*, an inhibitor of *IL1A/B*, was up-regulated in the tumor core (**Fig. 5C**). *IL1RN* is an important anti-inflammatory regulator that acts as a dominant negative interactor with *IL1R1* to actively suppress immune activation. Another, highly enriched gene in immune cells within the tumor core was *TGFBI* (**Fig. 5C**) a molecule with a yet unclear role in tumor formation (Han et al., 2015), previously thought to be expressed by neoplastic cells. *TGFBI*, which is induced by TGF-beta, is part of the non-Smad mediated TGF-b signaling pathway. The expression of *TGFBI* by immune cells of the tumor microenvironment could be potentially beneficial for tumor growth and dissemination since *TGFBI* has been shown to inhibit cell adhesion (dissemination), promote survival of cells with DNA damage (survival/resistance to treatment) and is a potential angiogenic factor (growth). Immune cells within the tumor also seem to further promote angiogenesis via the expression of *VEGFA* (**Fig. 5C**), a hypoxia induced (via *HIF1A*) angiogenic factor that promotes both vascular permeability and endothelial cell growth.

We also compared the expression of immune checkpoint receptor ligands in myeloid cells from the tumor core versus the peritumoral tissue. We found a small but statistically significant expression difference (p_adj_<0.01, as determined by DESeq2) between tumor and peritumoral myeloid cells for both ligands of the receptors PD1 (*CD274* (*PDL1*), *PDCD1LG2* (*PDL2*)) and *CTLA4* (*CD80*, *CD86*), as well as genes *ICOSLG* (ligand of ICOS receptor), *CD276* (B7-H3), *TNFRSF14* (ligand of *BTLA*) and *LGALS9* (ligand of *TIM3*). All genes were found to be up-regulated in the peritumoral compartment, with the exception of *LGALS9* and *CD80* that were up-regulated in the tumor (**Fig. S14**). The fraction of immune cells expressing each of the above genes is shown in Table S5.

## Conclusions

Glioblastoma is the most frequent and deadly primary brain cancer in adults (Kleihues and Sobin, 2000). Current treatment strategies combining surgery with radiotherapy and chemotherapy prolong survival, but tumor recurrence usually occurs within two years. Recurrence stems from infiltrating neoplastic cells originating initially from the tumor core, spreading quickly and across long distances within the brain. Thus, there is great hope in new treatment strategies that specifically target neoplastic cells in general as well as the infiltrating populations of cells migrating away from the primary tumor. In addition, novel immunotherapies are also being developed (Binder et al., 2016) relying on tumor vaccinations, immune checkpoint blockade, adoptive T-cell transfer and combinatorial immunotherapies.

This work represents the first identification and characterization of individual infiltrating tumor cells in the tissue surrounding the GBM tumor core. Furthermore, our high-depth and full-length RNAseq data allowed us to use gene expression information to infer genomic variation on the level of large chromosomal aberrations as well as smaller genomic variants such as insertions/deletions and single nucleotide somatic mutations. Our analysis revealed a large degree of heterogeneity between and within different tumors. We found that different patients’ tumors harbor different chromosomal aberrations while sharing hallmark CNVs such as the amplification of chromosome 7. Furthermore, each patient’s neoplastic cells have a unique set of smaller-scale mutations that also demarcate different populations of neoplastic cells within each tumor and potentially derive from different intra-tumoral lineages.

We defined infiltrating tumor cells as a population of neoplastic cells originating from the peripheral tissue whose transcriptional and genomic variant profiles resembled tumor core cells. Despite the heterogeneity of neoplastic cells originating from each of the individual patients, infiltrating cells share common characteristics regardless of patient of origin. The homogeneous gene signature of infiltrating GBM cells presents the potential of a convergent strategy for the mechanism of infiltration between highly variable tumors, which may provide new therapeutic avenues. When we examined the genes specifically up-regulated in infiltrating neoplastic cells, we found groups of genes involved in size regulation, energy production, inhibition of apoptosis, regulation of cell-cell adhesion as well as CNS development. In addition, infiltrating cells are likely hijacking machinery used during CNS development or later by cells migrating over long distances. It should be noted that the specific gene expression changes observed in infiltrating cells may either be intrinsic to the cell type or a result of migrating through a non-tumor microenvironment.

We also characterized the effect of the tumor microenvironment on non-neoplastic cells. Among the non-neoplastic cell populations, OPCs, neurons, mature oligodendrocytes, and vascular cells originating from both the tumor core and surrounding regions are essentially indistinguishable from each other In contrast, myeloid cells, consisting primarily of macrophages and microglia, are greatly affected by the tumor microenvironment. Despite belonging to the same cell class, myeloid cells within the tumor display unique gene expression profiles, demonstrating the direct effect of the tumor milieu on these important immune mediators. Specifically we found that immune cells within the tumor mass play a crucial role in enhancing tumor growth, survival and dissemination via suppression of inflammation, promotion of angiogenesis and ECM remodeling. Additionally, a subset of the changes induced by the tumor are shared between macrophages and microglia. These data, in addition to the full transcriptomes of immune cells within and surrounding the tumor, may help refine the development of novel therapies targeting cells that enhance tumor growth but are not themselves neoplastic. In particular, the findings from our study have implications for therapeutic immunotherapy efforts in GBM. For example, while we did find robust expression of MHC class I gene expression, which would allow for T-cell mediated immune responses, the expression of these genes varied greatly between patients. We also observed similar expression patterns for the ligands of PD1, which play a role in abrogating immune responses in some tumors. These data suggest that targeted immune therapies may vary vastly between patients, and that phenotyping patient tumors either via tumor resection or using technologies that capture tumor cells non-invasively in the CSF may be essential for screening patients who may be particular responsive to therapy.

## Author contributions

S.D., S.A.S., M.H.G. and S.R.Q. designed research; S.D., S.A.S., C.C., M.H.G., N.N. and M.K. performed research; M.H.G., G.L., S.C., I.C. and Y.L. contributed new reagents/analytic tools; S.D., D.C., S.A.S., and S.R.Q. analyzed data; S.D., S.A.S. and S.R.Q. wrote the paper.

## Acknowledgments

The authors thank Jennifer Okamoto and Gary Mantalas for assistance with sequencing and library preparation. This study was supported by The Hearst Foundation, and The Curci Foundation, NIH Grants K08-NS901527, R21-CA193046-01 (to M.H.G), by California Institute for Regenerative Medicine Grant GC1R-06673-A, Center of Excellence for Stem Cell Genomics (to S.R.Q.), and by the US National Institute of Health (NIH) R01 MH099555-03 (to B.A.B.). S.D. was supported by a fellowship from Svenska Sällskapet för Medicinsk Forskning. SAS was supported by NIMH T32GM007365, F30MH106261 and Bio-X Predoctoral Fellowships.

## Supplemental figure and table titles and legends

**Supplementary Figure 1.** Quality control of all single cells using houskeeping genes. Cells with high expression of housekeeping genes (light blue colored cluster) are selected for downstream analysis.

**Supplementary Figure 2.** Histograms of EGFR (A) and SOX9 (B) for regular and neoplastic cells. ROC performance analysis of EGFR (C) and SOX9 (D) expression as a classifier for neoplastic and regular cells and combined performance if both genes are used (E).

**Supplementary Figure 3.** TSNE showing aggregated single cell data from three datasets (GBM, Healthy brain and Bulk GBM). Single cells are colored by dataset (A), neoplastic or regular (B) and by cell-type (C).

**Supplementary Figure 4.** Boxplots showing values of “hypoxia-score”, ie normalized sum of expression for genes PGK1, CA9, VEGFA, SPP1, HIF1A for each neoplastic and regular cell, grouped by location (peripheral tissue or tumor core). Higher values denote cells at more hypoxic environments. Neoplastic cells from the periphery are the infiltrating neoplastic cells. All populations are different in a statistical significant manner (p<0.01, by a Wilcoxon rank sum test, with multiple testig correction) with the exception of neoplastic and regular cells from the periphery where p<0.05.

**Supplementary Figure 5.** A. Differentially expressed genes between neoplastic and non-neoplastic cells. For each gene the fraction of expressing neoplastic and non-neoplastic cells is plotted. B. Fraction of non-neoplastic cells, separated by cell type, expressing each of the neoplastic-enriched genes in FigS4A.

**Supplementary Figure 6.** *in situ* RNA staining for ALDH1L1 and MBP on tumor core and peripheral tissue.

**Supplementary Figure 7.** Boxplots showing expression of MHC I genes (A) and genes coding for ligands of PD1 (CD274 and PDCD1LG2) and CTLA4 (CD80 and CD86) (B) on neoplastic cells per patient. (C) Co-expression of CD274 and PDCD1LG2 across neoplastic cells of all patients. Units of all numerical axes are log2 CPM.

**Supplementary Figure 8.** (A) TSNE zoom in showing neoplastic cells colored by patient. (B) Heatmap showing expression of differentially expressed genes in neoplastic cells from each patient. (C) Histogram distributions of all cell-to-cell pairwise distances for each of the patient specific neoplastic clusters (shades of orange) and clusters of healthy non-neoplastic cell populations (shades of green).

**Supplementary Figure 9.** RNAseq based CNV analysis of neoplastic and non-neoplastic cells. Upper panel, individual median CNV profiles of neoplastic and non-neoplastic cells. Middle panel, difference in mean CNV profiles between neoplastic cells and non-neoplastic, non-myeloid cells. Regions of chromosome that are potentially amplified are pseudocolored green. Potential deletions are pseudocolored red. Lower panel, counts of positive and negative occurences of the DeltaCNV profile for each chromosome

**Supplementary Figure 10.** CNV analysis of DNAseq derived data for patient BT_S4. Distibution of the ratio of genomic reads between DNA from the tumor and from the peripheral cortex across the whole genome with the exception of chromosome Y. The ratio is shown on the right y-axis while estimated ploidy on the left.

**Supplementary Figure 11.** Presence of tumor specific variants shown in Figure 3C in non-neoplastic cells.

**Supplementary Figure 12.** GO analysis of genes up-regulated in infiltrating neoplastic cells compared to core neoplastic cells.

**Supplementary Figure 13.** PAGODA analysis of infiltrating (n=60) and core tumor (n=60) neoplastic cells. Significant aspects of heterogneity annotated by GO are shown.

**Supplementary Figure 14.** Boxplots showing expression in immune cells from the tumor and periphery of differentially expressed genes (padj<0.01, as determined by DESeq2) involved in immune-checkpoint regulation.

**Supplementary Figure 15.** permanent pathology H&E specimens showing representative tumor core, peripheral brain, and the tumors’ infiltrating margin for each patient included in the study.

**Table S1.** Top20 most enriched genes in each single-cell cluster. Color and naming of each cluster is consistent with Figure 2A.

**Table S2.** Number of cells per tSNE cluster and anatomical location along with each cluster’s cell identity.

**Table S3.** Classification of cells in the Healthy brain dataset classified to major cell types of the brain using the GBM dataset as reference.

**Table S4.** Fraction of neoplastic cells per patient expressing each of the genes shown.

**Table S5.** Fraction of immune cells separated by tissue of origin expressing each of the immune checkpoint genes shown.

**Table S6.** Classification of neoplastic cells to each of the four GBM subtypes.

**Table S7.** Primary and secondary antibodies used in the study.

**Table S8.** Sequences of oligonucleotides used for in situ RNA staining.

**Table S9.** Clinical information of patients included in the study. WT: Wildtype, M: Methylated, NM: Not Methylated, NT: Not tested.

## CONTACT FOR REAGENT AND RESOURCE SHARING

Further information and requests for resources and reagents should be directed to and will be fulfilled by the Lead Contact, Stephen R. Quake

## EXPERIMENTAL MODEL AND SUBJECT DETAILS

Clinical information for each patient included in the study can be found on **Table S9**. Informed consent was obtained from all subjects.

## METHOD DETAILS

### Tissue dissociation, immunopanning and single cell sorting

The experimental layout is outlined in **Fig. 1B**. We analyzed samples from four patients with confirmed cases of primary GBM. An additional sample from each patient was sent to pathology to confirm the diagnosis of GBM and to identify IDH1 status (negative). From each patient we collected two separate tissue samples, one originating from the tumor core and another from the peritumoral space (cortex) immediately adjacent to the tumor core. The tumor core was demarcated on MRI as strongly enhancing (with gadolinium contrast) (**Fig. 1A**) unlike the non-contrast enhancing peritumor space. Peritumor cortex was always removed prior to resecting tumor core in order to prevent potential cross-contamination. For each sample, tumor and peritumor tissue were processed separately.

Tissue samples were transported to the laboratory immediately from the operating room in order to begin dissociating samples within 1 hour of resection. The samples were processed similarly to existing protocols for human brain dissection^9^. Briefly, the tissue was first chopped into small pieces <1mm^3^ using a #10 scalpel blade and then incubated in 30 unit/ml papain at 34°C for 100 minutes. After digestion, the tissue was washed with a protease inhibitor stock solution. The tissue was then gently triturated in order to yield a single cell suspension. The single cell suspension was then added to a series of plastic petri dishes pre-coated with cell type specific antibodies (see below) and incubated for 10 - 30 minutes at room temperature. Unbound cells were transferred to the subsequent petri dish while the dish with bound cells was rinsed 8 times with PBS to wash away loosely bound contaminating cell types. The antibodies used include anti-CD45 to capture microglia/macrophages, anti-O4 hybridoma to harvest oligodendrocytes lineage cells, anti-Thy1 (CD90) to harvest neurons, anti-HepaCAM to harvest astrocytes, and *Banderiaea simplicifolia* lectin 1 (BSL-1) to harvest endothelial cells. Once bound to the Petri dish and rinsed, adherent cells were removed by incubating in a trypsin solution at 37°C for 5-10 minutes before gently squirting the cells off the plate. These single cell suspensions were then transferred to a FACS buffer before proceeding with single cell sorting.

Single cell suspensions were sorted using a SONY SH800 Cell Sorter. All events were gated using four consecutive gates: i) FCS-A/SSC-A, ii) FCS-H/FCS-W, iii) Propidium Iodide (PI) negative and iv) Hoechst positive. Single-cells were sorted in 96-well plates containing 4ul of lysis buffer (4U Recombinant RNase Inhibitor (TAKARA BIO), 0.05% Triton^TM^ X-100 (Thermo Fisher), 2.5mM dNTP mix (Thermo Fisher), 2.5uM Oligo-dT_30_VN (5′–AAGCAGTGGTATCAACGCAGAGTACT_30_VN-3′)), spun down for 2 minutes at 1000g and snap frozen. Plates containing sorted cells were stored at −80°C until processed.

### cDNA synthesis and library preparation

Reverse transcription (RT) and PCR amplification was performed using the Smart-seq2 protocol, described in (Picelli et al., 2014). Briefly, 96-well plates containing single-cell lysates were thawed on ice followed by incubation at 72 °C for 3 minutes and placed immediately on ice. Reverse transcription was carried out after adding 6ul of RT-mix (100U SMARTScribe^TM^ Reverse Transcriptase (TAKARA BIO), 10U Recombinant RNase Inhibitor (TAKARA BIO), 1X First-Strand Buffer (TAKARA BIO), 8.5mM DTT (Invitrogen), 0.4mM Betaine (Sigma), 10mM MgCl_2_ (Sigma) and 1.6uM TSO (5′-AAGCAGTGGTATCAACGCAGAGTACATrGrG+G-3)), for 90 minutes at 42°C, followed by 5 minutes at 70°C.

RT was followed by PCR amplification. PCR was performed using 15 ul of PCR-mix (1x KAPA HiFi HotStart ReadyMix (Kapa Biosystems), 0.16uM ISPCR oligo (5′-AAGCAGTGGTATCAACGCAGAGT-3′ of Lambda Exonuclease (NEB) using the following thermal-cycling protocol: 1. 37°C for 30min, 2. 95°C for 3min, 3. 21 cycles of 98°C for 20s, 67°C for 15s and 72°C for 4 min and 4. 72°C for 5 min.

PCR was followed by bead purification using 0.7x AMPure beads (Beckman Coulter), capillary electrophoresis and smear analysis using a Fragment Analyzer^TM^ (AATI). Calculated smear concentrations within the size range of 500 and 5000 bp for each single cell were used to dilute samples for Nextera library preparation as described in (Darmanis et al., 2015).

### Sequencing and QC

In total, we sequenced 4028 single cells using 75bp-long paired-end reads on a NextSeq instrument (Illumina) using High-output v2 kits (Illumina). Raw reads were preprocessed and aligned to the human genome (hg19) using the exact pipeline described in (Darmanis et al., 2015).

As a quality metric, we first performed hierarchical clustering on all cells using a list of housekeeping genes (Fig. S1), and removed any cells with uniformly low expression across all genes (likely a result of low quality RNA or cDNA synthesis). We separated the resulting dendrogram into two clusters containing cells that passed (n=3589) or failed (n=456) this quality control (QC). All downstream analyses were performed using only the cells that passed QC. A summary of the number of cells by QC cluster (pass or fail), with respect to patient and tissue of origin along with sequencing statistics can be found in **Table 1.**

### Preparation of libraries from genomic DNA

Human genomic DNA was sheared to an average size of 500bp using a Covaris S220 following the manufacturer’s protocol in snap cap microtubes. Sheared DNA (500ng) was ligated into indexed Illumina sequencing adapters (made by IDT) with the Kapa Hyper Prep Kit for Illumina. Libraries were quantitated using average DNA fragment length (AATI Fragment Analyzer) and concentration determined by qPCR (Kapa Library Quantitifcation Kit for NGS). Pooled libraries were sequenced as paired end 150bp reads on an Illumina NextSeq500.

### Antibody staining

Tissue sections were removed from −80^0^C and allowed to dry at RT for 30mins followed by fixation with PFA (4%, in PBS). Sections were washed three times using PBS (5 min washes at RT) and blocked with blocking buffer (1xPBS, 10% donkey serum, 0.1% Triton) for 1h at RT. After blocking sections were probed with primary antibodies (for antibodies used and working concentrations see **Extended data Table 7**) and incubated overnight at 4^0^C. Sections were washed and secondary antibodies (Table SX) were added and incubated with the tissue for 1h at 37^0^C followed by three 5 minute washes with PBS. Sections were stained with DAPI (1x in PBS) for 30min at RT, washed another two times as before, dried and mounted with SlowFade Gold antifade reagent (Life Technologies) prior to imaging.

### In situ RNA

In situ RNA staining was performed on 10 µm thick fresh-frozen tissue sections using padlock probes and rolling-circle amplification (RCA) as described before (Ke et al., 2013). In situ reverse transcription was carried out with primers containing 2-O-Me modified bases. The sequence of the primers, padlock probes and fluorescently labeled detection oligos are listed in extended data table 8. In order to reduce the high fluorescence background due to the presence of lipofuscins in the biopsy from tumor periphery, the sections were incubated in 1% Sudan Black in 70% Ethanol for 30 minutes at RT in the dark prior fluorescence labeling of RCA products.

Multiplexed detection was achieved via combinatorial labeling of RCA products by sequential hybridization four distinct fluorescence detection oligos. Three cycles of hybridization, imaging and detection oligo removal allow identification of 64 gene-specific fluorescence barcodes (43) in principle enough to distinguish 54 probes used in this study. To further reduce the complexity of the barcodes and facilitate the identification of false positive (errors), we split the padlock probes in two distinct pools of 27 and 27 probes and performed the in situ RNA staining on consecutive sections. Also, to distinguish true signals from tissue auto fluorescence (as described below) only two colors are used to label RCA products at third cycle.

## QUANTIFICATION AND STATISTICAL ANALYSIS

All data analysis was performed using R. Specific packages and functions used are described in greater detail below. All code is available upon request.

### Dimensionality reduction and clustering

Dimensionality reduction was performed in three steps. First, we calculated the overdispersion of each gene as described (Fan et al., 2016). We then selected the top 500 over-dispersed genes and constructed a cell-to-cell distance matrix (1-absolute correlation) of all cells. The distance matrix was reduced to two dimensions using tSNE as previously described in (Darmanis et al., 2015) and as implemented in package “tsne” for R (perplexity=50). Clustering of groups of similar cells was performed on the 2-dimensional tSNE space using kmeans as implemented in package “stats” for R.

### CNV analysis

We constructed CNV vectors for each single cell based on gene expression data. Given the nature of RNAseq data, CNV profiles cannot be calculated using the same approach as when genomic DNA data are available. Instead, one can use the gene expression information to infer over- or under-expression of big genomic regions that might correspond to chromosomal amplification or deletion events. To calculate CNV profiles for each single cell we used a similar approach to (Patel et al., 2014) and (Tirosh et al., 2016). Briefly, we sorted all genes based on their genomic location and calculated a CNV vector for every cell. The CNV vector is a moving average of gene expression using a window of 0.1*n genes per chromosome, where n is the total number of genes on that chromosome. The resulting CNV vectors of each cell were centered by subtraction of their mean prior to any downstream analysis.

### Differential expression analysis

Differential expression analysis was performed using package DESeq2 (Love et al., 2014) for R. All functions in the DESeq2 pipeline were used with default settings.

### Differential expression analysis between neoplastic and non-neoplastic cells

Using DESeq2, we found a large number of genes (n_total_=5143, n_upregulated_=3985, n_downregulated_=1158) that were either up- or down-regulated in neoplastic cells compared to non-neoplastic cells of the brain. To limit the list of differentially expressed genes to those that showed specific tumor-restricted expression we looked for genes that were expressed (expression was strictly defined as the presence of even a single read for each gene) in more than 60% of all neoplastic and in less than 20% of all non-neoplastic cells. The percentage of expressing neoplastic and not-neoplastic cells for each of the genes that fulfilled our criteria (n=30), along with the type of healthy cells expressing each gene are shown in **Figure 2A.** We also noted that the majority (~50% on average) of non-neoplastic cells expressing genes up-regulated in neoplastic cells belong to the OPC cluster (**Figure 2B**).

### Verhaak classification analysis of neoplastic cells

We initially thought that the differences between patients could be attributed to the fact that each tumor belongs to a different GBM subtype, namely one of the Proneural (PN), Neural (NL), Mesenchymal (MES) and Classical (CL) subtypes *(4)*. To test this hypothesis, we used 792 genes from the list of 800 genes that were used in *(4)* to classify different GBM samples to the four molecular subtypes.

Using the list of 792 genes, we calculated the correlation between each of the neoplastic single cells and an average expression profile for each of the four GBM subtypes. We then assigned each cell to the subtype with the highest correlation. We found that the majority of cells within each patient were classified as CL with the remainder of the cells belonging to the remaining categories, as summarized in **Extended data Table 6.** In our opinion, based on the classification of each patient’s cells, different GBM subtypes do not seem to be responsible for the observed differences between patients. Nonetheless, neoplastic cells originating from different tumors are fundamentally different both in terms of their gene expression profiles, as can be seen in **Figure S8**, as well as their individual CNV profiles shown in **Figure 3**.

### Myeloid cell identity

We inferred the identity of each cell in the two myeloid clusters using ImmGen’s (https://www.immgen.org/) microarray gene-expression data from 214 purified mouse immune cell populations belonging to 22 different broad classes of immune and supporting cell types. We correlated each single cell in our myeloid clusters to each of the 214 ImmGen samples. We then selected the top 5 correlated ImmGen samples for each single cell and assigned an immune-cell type to each single cell using a majority vote.

### Genomic DNA data analysis

Raw DNA sequences were trimmed for low quality bases and Illumina adapter sequences using Trimmomatic (version 0.32, default parameters). Paired end reads from fragments shorter than twice the read length were merged and a consensus sequence of the overlapping bases was determined using FLASH (version 1.2.11). Reads were then aligned to the UniVec core database ((Cancer Genome Atlas Research Network, 2008; Verhaak et al., 2010)) using bowtie2 (version 2.2.4, –local mode) to remove sequences derived from common vector/control sequences, including the bacteriophage phi X 174, a spike in control used with Illumina sequencers. The final set of reads was then aligned using bowtie2 to the human reference (GRCh38), with the results saved as a BAM file. Each sample was sequenced to an average depth of ~20X.

Resulting BAM files were analyzed for the presence of chromosomal aberrations using the R package CNAnorm. No GC correction was used.

### PAGODA analysis

We performed a similar analysis using PAGODA (Patel et al., 2014) in an attempt to group genes responsible for differentiating between infiltrating and core tumor cells, in groups of similar biological functions. Due to a very long processing time when all cells are used, we performed the analysis using all infiltrating cells and only an equal number of randomly selected core neoplastic cells. As a part of the analysis, we identified pre-defined genes sets (representing all GO terms) that exhibited statistically significant abundance of coordinated variability. All top aspects of heterogeneity, grouped by GO category, are shown in **Fig. S13.** To our satisfaction, we found that the genes underlying the top aspects of heterogeneity identified by PAGODA are very similar to the genes we identified with our prior analysis. The genes are also grouped in relevant GO categories such as response to external stimulus, tissue development, cell projection organization and glial cell proliferation.

### Image acquisition and analysis

Stained sections were imaged on a Zeiss Axioplan epifluorescence microscope equipped with filter-cubes for DAPI, FITC, Cy3, TexasRed and Cy5, an Axiocam 506 mono camera (Zeiss), automated filter-cube wheel and a motorized stage. Z-stacks of 12 images were acquired with a 20X (0.8NA) Plan-Apochromat objective and maximum intensity projections (MIP) were generated with Zen 2.3 image acquisition software. Retrospective illumination correction was performed using CellProfiler as described in Singh et al (Singh et al., 2014). The generated illumination correction function was then used as reference in the shade correction module of Zen 2.3 software.

Images were pre-processed using FIJI software. Briefly, MIP images were cropped and aligned based on DAPI nuclear staining using rigid-transformation function in the MultiStackReg plugin. Then a mask of the RNA staining containing all the RCA products was created by combining the two single channels images coding for the last staining cycle in each pool. The remaining two channels were removed from the mask in order to attenuate background fluorescence from lipofuscins in the brain which is visible in all the fluorescence spectra used. Pre-processed images were then loaded in Cellprofiler for spot detection and intensity measurements and nuclei counts used to normalize the in situ RNA data. Barcode identification was carried out using the same Matlab script described here (Ke et al., 2013). All script and images are available upon request. A quality threshold was applied to the barcode calling until the number of non-expected barcodes from each image was around 1% (max 1.3, min 0.1 for four image sets). Homo-polymer barcodes identify background fluorescence and were removed. After quality thresholding, genes with lower counts than the highest non-expected barcode were considered not detected. The counts of each detected barcode were normalized by the number of nuclei in the corresponding section in order to compare gene expression among the different samples.

### Variant analysis

Single cell variants were called using the GATK pipeline for single cell RNAseq data using the suggested arguments. Reads were first aligned to the hg19 human reference using STAR (version 2.4.2a run in two-pass mode), following which we used the GATK tools MarkDuplicates, SplitNCigarReads, BaseRecalibrator, and HaplotypeCaller. Potential variants were then filtered using VariantFilter (-window 35 -cluster 3 -filterName FS -filter "FS > 30.0" -filterName QD -filter "QD < 2.0"). The SNPiR pipeline (Piskol et al., 2013) was employed to further exclude variants belonging to known RNA editing sites, variants in intronic positions near splice junctions, as well as variants in non-uniquely mapped, repetitive, and homopolymer regions. Lastly, variants were annotated using ANNOVAR (Wang et al., 2010) and retained only if observed in at least three cells.

To confirm abundant RNAseq tumor variants in patient BT_S4, we used the Varscan 2 *somatic* tool (Koboldt et al., 2012) (version 2.3.9) on paired tumor-peripheral DNA whole genome sequencing data aligned to the human reference with BWA (version 0.7.15). To sensitively detect potentially rare variants while accounting for the substantial fraction of healthy cells in the tumor and the potential for infiltrating cells in the periphery, the following parameters were used: –min-coverage 5 –normal-purity 0.95 –tumor-purity 0.5 –min-var-freq 0.001.

## DATA AND SOFTWARE AVAILABILITY

The data reported in this paper have been deposited in the Gene Expression Omnibus (GEO) database, (accession no. GSE84465).

